# Two linked loci under mutation-selection balance and Muller’s ratchet

**DOI:** 10.1101/477489

**Authors:** Ksenia A. Khudyakova, Tatiana Yu. Neretina, Alexey S. Kondrashov

## Abstract

We report the complete analysis of a deterministic model of deleterious mutations and negative selection against them at two haploid loci without recombination. As long as mutation is a weaker force than selection, mutant alleles remain rare at the only stable equilibrium, and otherwise, a variety of dynamics are possible. If the mutation-free genotype is absent, generally the only stable equilibrium is the one that corresponds to fixation of the mutant allele at the locus where it is less deleterious. This result suggests that fixation of a deleterious allele that follows a click of the Muller’s ratchet is governed by natural selection, instead of random drift.

## Introduction

Negative selection against deleterious mutations occurs in all populations. As long as random drift can be ignored, such selection leads, with rare exceptions (Yang and Kondrashov, 2003), to deterministic mutation-selection equilibrium. Analysis of this equilibrium at a single locus represents one of the foundational results of population genetics (Haldane, 1937; Muller, 1950). Studies of a general multilocus case (Kimura and Maruyama, 1966; Hermisson et al., 2002) were based on an unrealistic assumption that mutations at all the loci are equally deleterious.

We relax this assumption in the simplest possible case of two totally linked loci in an infinite haploid population. This model is simple enough to be analyzed completely, but still rich enough to admit a number of qualitetively different regimes. The mode of epistasis between the two loci, synergistic vs. diminishing returns, is the key factor affecting dynamics of the population. Our results may be relevant to the process of fixation of a deleterious allele that follows a click of the Muller’s ratchet (Charlesworth and Charlesworth, 1997).

## Model

Consider two non-recombining haploid diallelic loci, A and B, where allele *A* mutates into a deleterious allele *a* with the rate μ and allele *B* mutates into a deleterious allele *b* with the rate ν. Frequencies of genotypes of *AB*, *Ab*, *aB*, and *ab* are 1-x-y-z, x, y, and z, respectively, and their fitnesses are 1, 1-t, 1-s, and (1-s)*(1-t)*(1-k), respectively, where s and t are coefficients of selection against mutant alleles at loci A and B, and k characterizes epistasis. Then, assuming discrete generations, genotype frequencies in successive generations are connected by the following system of difference equations:

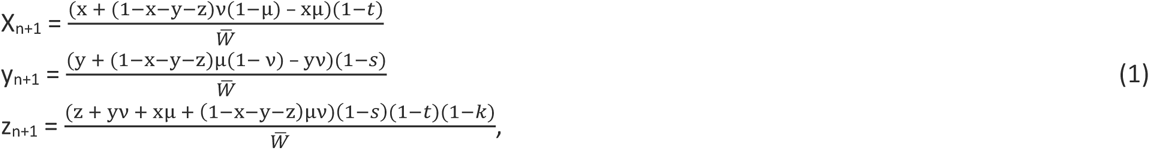

where

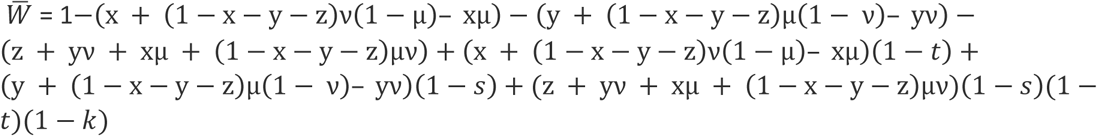

is the mean population fitness. Equilibria of this system and their stability were studied using Wolfram Mathematica.

## Results

### General case

Generally, (1) can have the following four equilibria (Fig. 1):

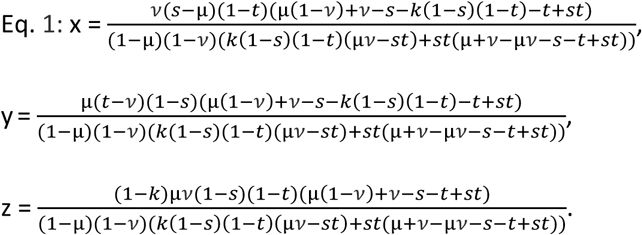

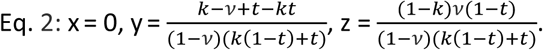

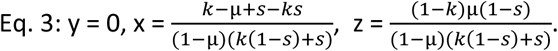

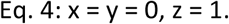

Also, if

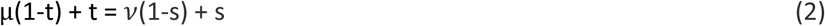

there is a straight line of equilibria which connects Eq. 2 and Eq. 3.

**Fig. 1.**
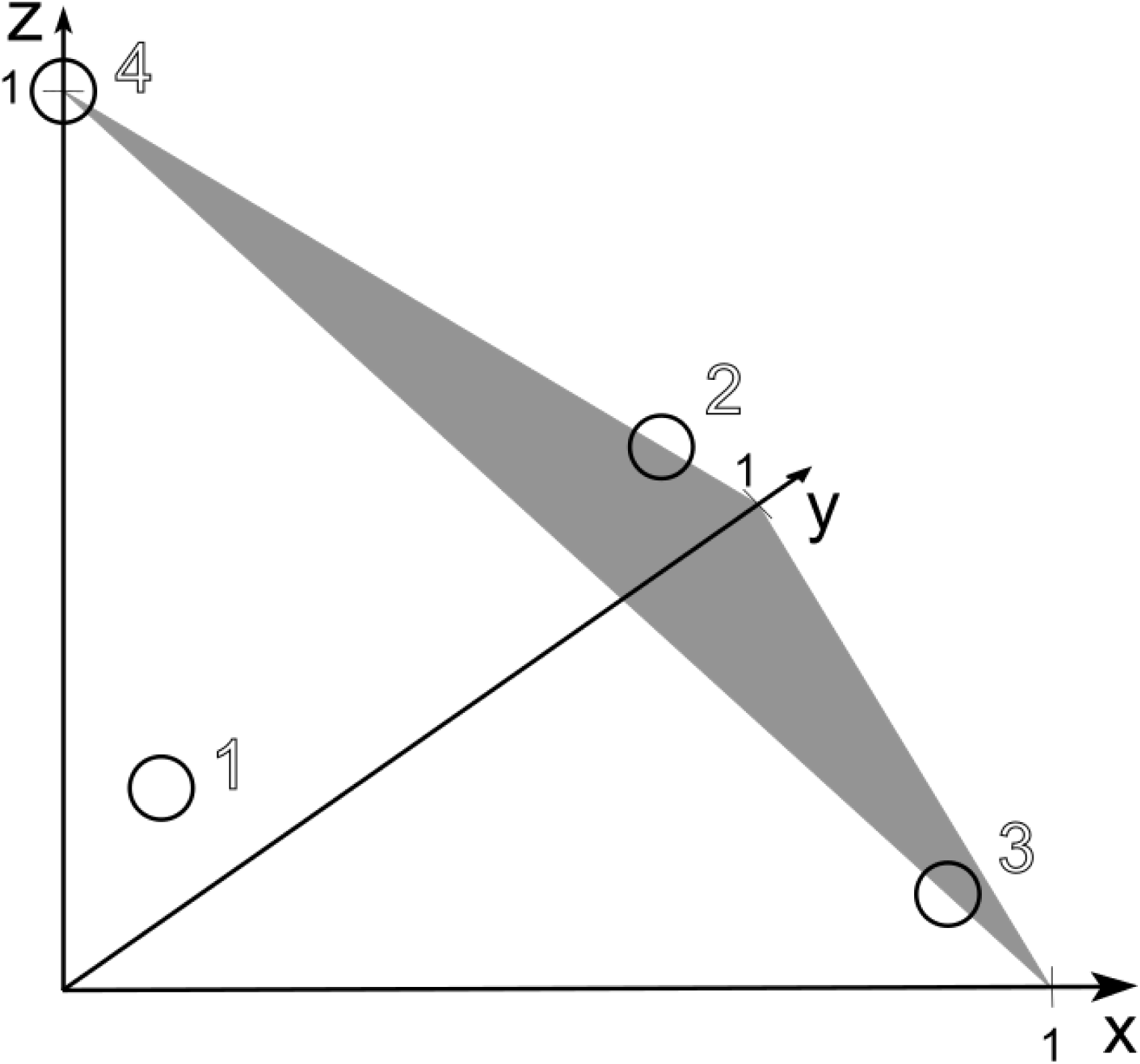
The four equilibria of system (1).

Eq. 1 describes the mutation-selection balance under which both wild-type alleles *A* and *B* are present, despite the mutational pressure. Eqs. 2 and 3 correspond to fixation of mutant allele *a* (Eq. 2) or *b* (Eq. 3) and, thus, describe the mutation-selection balance only at locus B (Eq. 2) or at locus A (Eq. 3). Eq. 4 corresponds to the fixation of mutant alleles at both loci.

When mutation is a weaker force than selection (μ << s, ν << t) Eq. 1 is the only stable one, and the corresponding frequencies of alleles *a* and *b* are low. Fig. 2 presents partitioning of the space of parameters μ, ν, s, t, and k into regions of qualitatively different dynamics. A case of s > t is shown, so that the straight line (2) intersects the abscissa axis when μ > 0. There are 6 and 7 such regions with synergistic (k > 0) and diminishing returns (k < 0) epistasis, respectively. To understand these patterns, it is helpful to keep in mind that under synergistic epistasis, negative selection provides a stronger protection to the wild-type allele at a locus when the other locus harbors the mutant allele than when it harbors the wild-type allele. By contrast, under diminishing returns epistasis, negative selection provides a weaker protection to the wild-type allele at a locus when the other locus harbors the mutant allele than when it harbors the wild-type allele.

Let us first consider synergistic epistasis (Fig. 2, top). Under ν < t, the following bifurcations happen when μ increases. When μ becomes larger than s, Eq. 1 merges with Eq. 2 which becomes stable. In this case, allele *A* always disappears from the population as long as allele *B* is present. When μ continues to increase and becomes larger than s+k(1-s), Eq. 3 merges with Eq. 4, which means that allele *A* disappears from the population even if allele *b* is fixed in it.

Under t < ν < t+k(1-t), when μ is small, there are three equilibria (Eq. 2, Eq. 3, and Eq. 4) and Eq. 3 is the only stable one. When μ increases so that μ(1-t)+t becomes larger than ν(1-s)+s, Eq. 3 and Eq. 2 switch their stabilities through a line of neutral equilibria, and Eq. 2 becomes stable. When μ becomes larger than s+k(1-s), Eq. 3 merges with Eq. 4.

Finally, under ν > t+k(1-t), when μ is small, only Eq. 3 and Eq. 4 are present, and Eq. 3 is stable. When μ becomes larger than s+k(1-s), Eq. 3 merges with Eq. 4 which becomes stable, which implies that mutant alleles *a* and *b* always reach fixation. Of course, analogous bifurcations occur when ν increases, under different values of μ.

Now let us consider diminishing returns epistasis (Fig. 2, bottom). Under ν < t+k(1-t), when μ becomes larger than s+k(1-s) Eq. 3 merges with Eq. 4. In this case, allele *A* always disappears from the population as long as allele *B* is absent. When μ continues to grow and exceeds s, Eq. 1 switches the stability with Eq. 2, which means that allele *A* disappears from the population even if allele *B* is present.

Under t + k(1-t) < ν < t, when μ is small, there are three equilibria (Eq. 1, Eq. 3, and Eq. 4) and Eq. 1 is the only stable one. When μ becomes larger than s+k(1-s), Eq. 3 merges with Eq. 4. When μ increases so that (μ-s)/(1-s) – k/2 becomes larger than -(ν-t)/(1-t) + k/2, Eq. 1 and Eq. 4 merge and switch their stabilities, so that alleles *a* and *b* always reach fixation.

Finally, under ν > t, when μ is small, there are only Eq. 3, which is stable, and Eq. 4. When μ becomes larger than s + k(1-s) Eq. 3 merges with Eq. 4 and switches the stability with it, which implies that mutant alleles *a* and *b* always reach fixation. Analogous bifurcations occur when ν increases, under different values of μ.

**Fig. 2.**
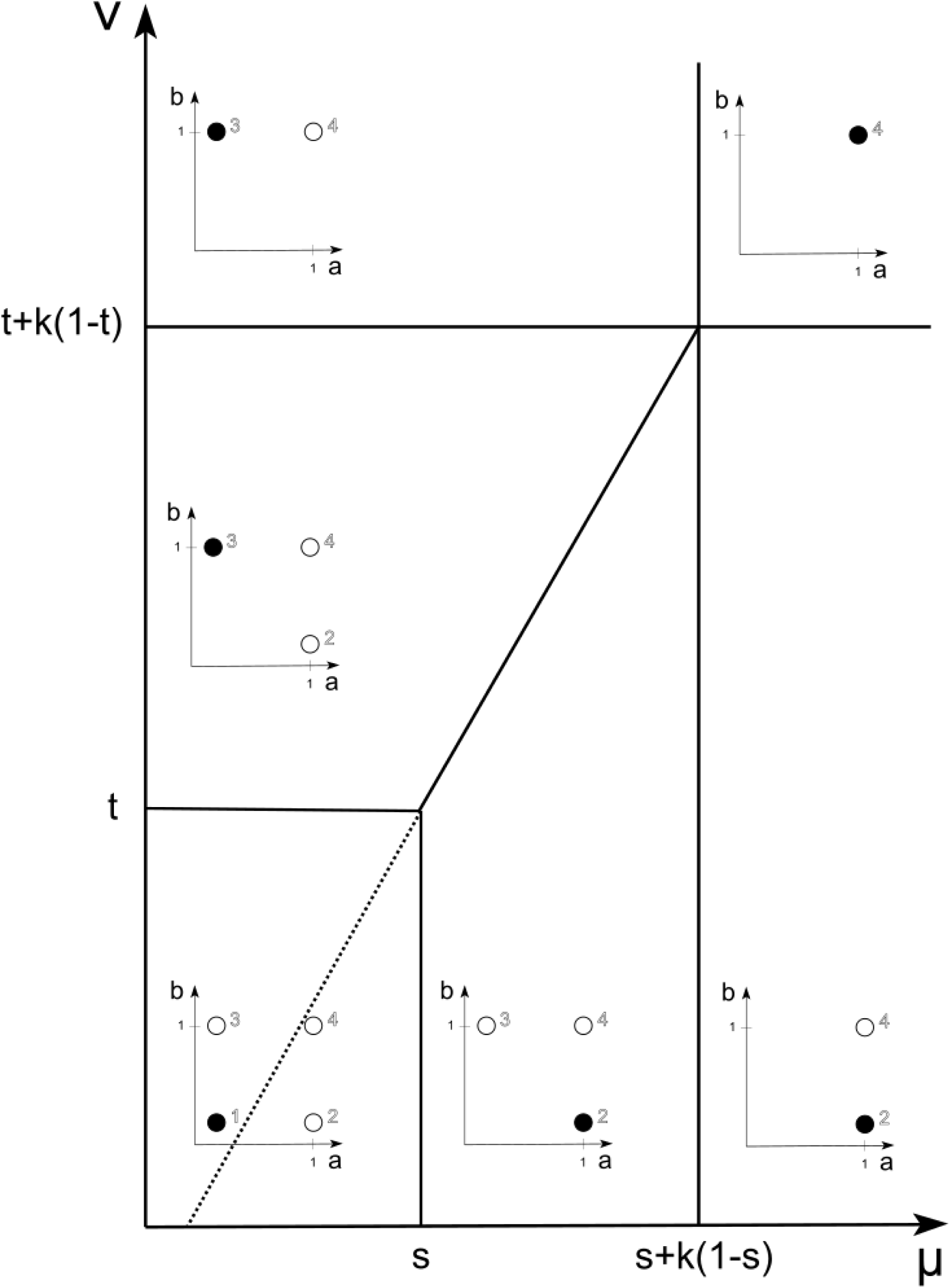
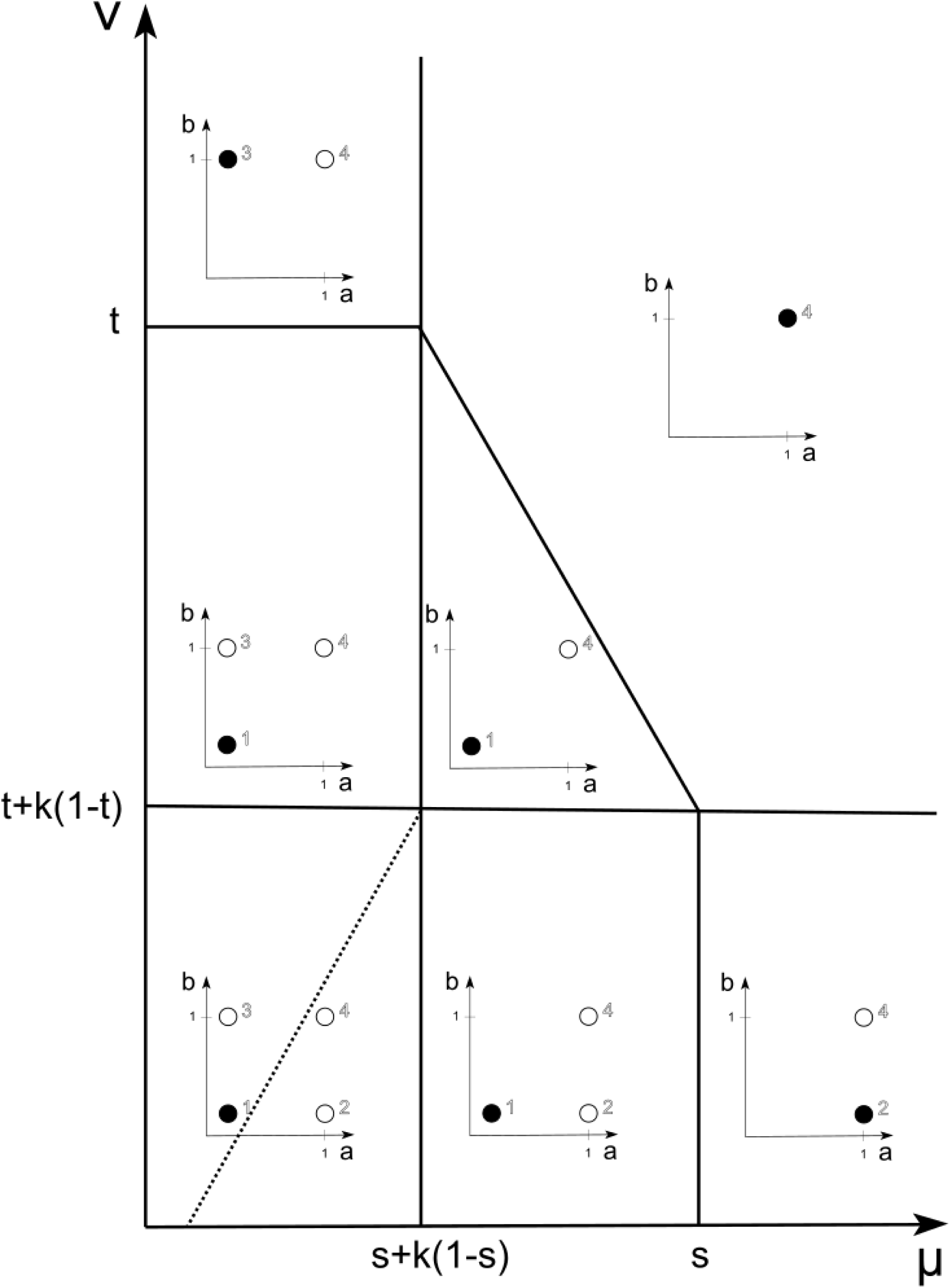
Qualitatively different dynamics of the system (1) under different values of parameters under synergistic (top) and diminishing returns (bottom) epistasis. Stable and unstable equilibria are shown as filled and empty circles, respectively. Instead of the 3-dimensional space of genotype frequencies, the locations of equilibria are shown on the plane that corresponds to different frequencies of alleles a and b. A click of Muller’s ratchet

Let us assume that μ < s and ν < t in the case of synergistic epistasis, or μ < s +k(1-s) and ν < t +k(1-t) in the case of diminishing returns epistasis, so that mutation cannot overwhelm selection at either locus. What happens if genotype *AB* is nevertheless absent from the population, perhaps due so its random loss, so that x+y+z = 1? In this case, dynamics of (1) are confined to a shadowed plane in Fig. 1 (Fig. 3).

**Fig. 3.**
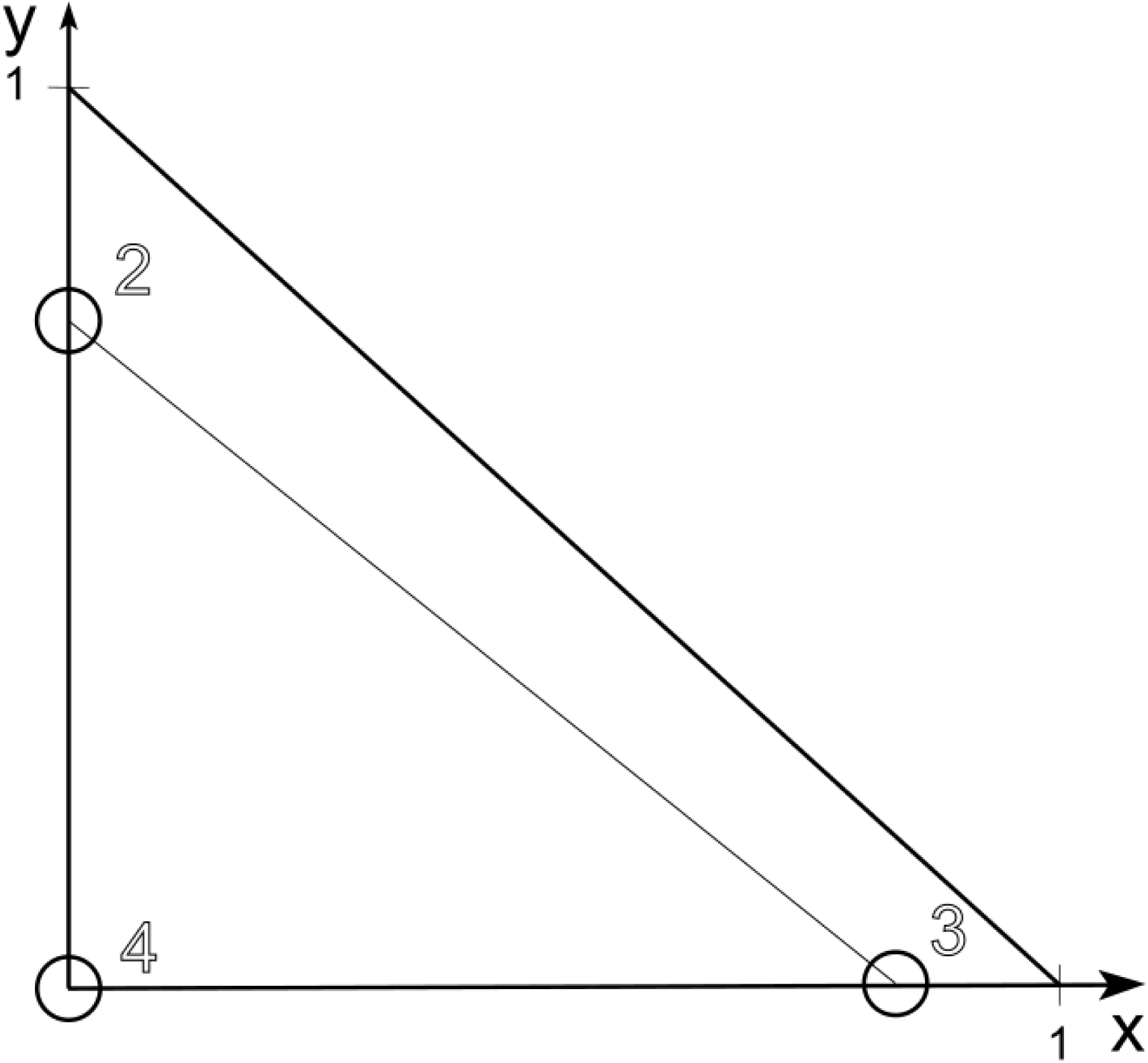
Three equilibria of system (1) in the case of a loss of genotype *AB* (shadowed plane on fig.1)

Within this plane, Eq. 2 is stable if the left-hand side of (2) exceeds the right-hand side, and Eq. 3 is stable in the opposite case. If (2) is an equality, Eq. 2 and Eq. 3 are connected by an attracting line of neutral equilibria. This is always the case, in particular, in the symmetric case, when both mutation rate and selection coefficients at the two loci are equal.

## Discussion

Consideration of the mutation-selection balance at two linked loci involves phenomena of two kinds. First, when mutation becomes a stronger force than negative selection, the loss of wild-type alleles can proceed through a variety of mechanisms, depending, in particular, on the mode of epistasis (Fig. 2). However, in this situation a deterministic model is likely to be unrealistic, because random drift also needs to be taken into account. Indeed, per locus mutation rates are of the order of 10^−5^ (if our loci are genes) or even 10^−8^ (if our loci are nucleotide sites). Clearly, when selection coefficients are below 10^−5^ or even 10^−8^, random drift becomes important in a population of any realistic size.

Second, in the absence of the mutation-free genotype, which can be thought of as a click of Muller’s ratchet, the mutant allele becomes fixed in one of the two loci (Fig. 3). This effect was discovered by Charllesworth and Charlesworth (1997), who emphasized random drift as the cause of this ratchet-caused fixation (see also Gordo and Charlesworth, 2001; Sӧderberg and Berg, 2007; Stephan and Kim, 2002; Stephan, Charlesworth and McVean, 1999; Gessler and Xu, 1999; Gordo and Charlesworth, 2000a,b). However, Charlesworth and Charlesworth (1997) also observed that “if there is variation in the selection coecffients, … clearly mutations with weaker effects will be more likely to become fixed”. Our analysis show how this deterministic ratchet-induced fixation occurs. Random drift governs the process of fixation only if mutations are equally deleterious or, more precisely, if there is an exact balance between mutation rates and coefficients of selection at different loci (2). Otherwise, in a sufficiently large population, the mutant allele is fixed at a locus where the selection is weaker and the mutation rate is higher.

## Acknowledgements

This work was supported by the Russian Science Foundation grant N 16-14-10173.

